# T cell and peripheral blood parameters define progression of autoimmune disease in the IL-2Rα KO model

**DOI:** 10.1101/345512

**Authors:** Genevieve N. Mullins, Kristen M. Valentine, Mufadhal Al-Kuhlani, Dan Davini, Kirk D.C. Jensen, Katrina K. Hoyer

## Abstract

IL-2Rα is required to generate the high affinity receptor for IL-2, a cytokine important in immune proliferation, activation, and regulation. Mice deficient in IL-2Rα (IL-2Rα-KO) develop systemic autoimmune disease and die from severe anemia between 18-80 days of age. These mice develop kinetically differing autoimmune disease, with approximately a quarter dying by 21 days of age and half dying after 30 days. This research aims to define immune parameters that distinguish cohorts of mice that develop early- and late-stage autoimmune disease in the IL-2Rα-KO genetic background. To investigate these differences, we evaluated complete blood counts (CBC), antibody binding of RBCs, T cell numbers and activation, and hematopoietic progenitor changes, to assess the extent of peripheral autoimmune hemolytic anemia and bone marrow failure. Early onset disease correlated with anti-RBC antibodies and lower hematocrit on day 19. We also found that predicted late stage-disease IL-2Rα-KO mice have higher numbers of developing memory CD4 and CD8 T cells and reduced AIHA at early ages. The expansion of CD8 T cells seen in IL-2Rα-KO mice is driven by unimpaired IL-2 signaling which correlated with increased IL-2Rβ expression. Using a simple CBC we were able to predict disease kinetics to explore mechanisms underlying early and late disease.

## Introduction

Mice deficient in IL-2 (IL-2-KO) or its receptor (IL-2Rα-KO or IL-2Rβ-KO) develop spontaneous, multi-organ autoimmune disease, including autoimmune hemolytic anemia (AIHA), bone marrow failure (BMF), inflammatory bowel disease, and dacryoadenitis (1–7). Disease in these mice is driven primarily due to a lack of functional regulatory T cells (Tregs), as decreased IL-2 signaling leads to reduced survival and suppressive function of Tregs (4, 8–10). AIHA severity can be measured by RBC count, hemoglobin levels, hematocrit, and RBC-specific antibodies.

IL-2 is a cytokine important for the proliferation of activated T cells, and the survival and function of Tregs (4, 10–12). The receptor for IL-2 is comprised of three receptor chains - IL-2Rα (CD25), IL-2Rβ (CD122), and the common gamma chain (γ_c_; CD132) (10, 13). Binding of IL-2 to its receptor activates three main signaling pathways, including MAPK, PI3K, and JAK1/3-STAT5 pathways (12–14). These pathways promote cell survival and proliferation, as well as STAT5-mediated gene expression. IL-15, a common-gamma cytokine, also utilizes the same β and γ receptor chains, while IL-7 shares the γ chain (14, 15).

Signaling through IL-2 lowers the threshold of TCR signaling required to initiate proliferation in CD8 T cells, but not in CD4 T cells (16). In CD8 T cells, the strength of IL-2 signaling is important in the development of memory, with low signaling strength associated with the development of central memory rather than effector memory (15, 17). Due to the shared IL-2Rβ subunit, these differentiation signals occur in response to both IL-2 and IL-15 (15). In CD4 Tregs, IL-2 signaling stabilizes STAT5-mediated gene expression of FoxP3 to promote suppressive function, which does not occur in other CD4 T cell subsets (8, 13). IL-2 does not activate the PI3K pathway in Tregs, likely due to high expression of the phosphatase PTEN in these cells (13).

We set out to evaluate the mechanisms underlying the difference in disease kinetics in IL-2R - KO mice. Since it is known that IL-2Rα-KO T cells do not proliferate in response to low-dose IL-2 while WT T cells do (3), we explored potential differences in IL-2 signaling cascades. Before disease induction on day 19, we found that symptoms associated with AIHA were more severe in mice that develop early disease versus late disease, and the frequency of common lymphoid progenitors (CLP) and peripheral CD4 and CD8 T cells was increased in early disease. Further, IL-2Rα-KO T cells responded normally to high-dose IL-2 but had a reduced signaling capacity in response to low-dose IL-2.

## Materials and methods

### Mice

BALB/c IL-2Rα-KO or control littermate mice (wildtype and heterozygous; WT) were bred and maintained in our specific pathogen-free facility in an Animal Barrier Facility in accordance with the guidelines of the Laboratory Animal Resource Center of the University of California Merced and under approval by the Institutional Animal Care and Use Committee. Survival of IL-2R - KO and littermate controls was monitored from day 10 onward.

### Complete Blood Count (CBC)

Cardiac punctures or terminal eye bleeds were performed immediately following cervical dislocation, and blood collected in heparinized tubes (18). Blood samples used for predictions were collected from mice aged 19d via the submandibular vein. CBC were evaluated within 8 hr on a Hemavet 950 Veterinary Hematology System (Erba Diagnostics).

### RBC Ab detection

Abs bound to RBCs were detected using flow cytometry similar to previously described (19). RBCs were freshly isolated from mice by terminal bleed or submandibular vein puncture, washed three times in room-temperature PBS, and resuspended to 1% RBCs. 10 μ! of 1% RBCs were incubated with either anti-mouse IgM-FITC (1:100 dilution; on ice; Jackson ImmunoResearch) or anti-mouse IgG-FITC (1:50 dilution; at 37 °C; Jackson ImmunoResearch) for 20-30 min. The percentage of RBCs bound by Ab was determined by flow cytometry.

### Flow cytometry

Abs were purchased from eBioscience, unless otherwise indicated. Ab staining was performed in PBS/2%FBS (Omega Scientific) for 30 min at 4°C unless otherwise noted. Cell viability was determined using eFluor506 viability dye (eBioscience). For T cell activation, lymphocytes and splenocytes were stained with anti-CD8α (53-6.7), anti-CD4 (RM4-5), anti-CD62L (MEL-14), anti-CD44 (IM7), and, for exclusion of non-relevant cells, anti-CD45R (B220; RA3-6B2), anti-CD11b (MI/70), anti-CD11c (N418), and anti-Ly-6G (Gr-1; RB6-8C5). To evaluate hematopoietic progenitor populations in the BM, RBC-lysed BM cells were first incubated with biotinylated lineage markers - anti-CD3ε (145-2C11), anti-CD4 (GK1.5), anti-CD8α (53-6.7), anti-CD 11b (MI/70), anti-CD11c (N418), anti-Ly-6G (Gr-1; RB6-8C5), anti-CD45R (B220; RA3-6B2), and anti-Ter-119 (TER-119) - without Fc-block. Cells were washed, then stained anti-CD117 (c-kit; 2B8), anti-Ly6A/E (Sca-1; D7), anti-CD34 (RAM34), anti-CD16/32 (93), anti-CD127 (IL-7Rα; A7R34), and Streptavidin-APCe780. To evaluate RBC precursors, whole BM cells were incubated with anti-Ter119 (TER-119) and anti-CD71 (R17217). To assess proliferation, staining was performed as previously described (5). Cells were surface stained for 30 min at 4°C with anti-CD8α (53-6.7), anti-CD4 (RM4-5), biotinylated anti-CD3ε (145-2C11), and Annexin V (Invitrogen). Cells were then fixed using the FoxP3 Fixation Kit according to manufacturer’s directions, then stained for 45 min at room temperature for anti-Ki67 (SolA15), anti-FoxP3 (FJK-16s), and Streptavidin-APC. Flow cytometry was performed on a Becton Dickinson LSR-II and data analyzed using FCS Express with Diva Version 4 (DeNovo Software) or FlowJo.Version 7.6.5 (FlowJo).

### Intracellular Cytokine Stains

For evaluation of cytokine production, intracellular cytokine staining for IL-2 and IFNγ was performed as previously described (18, 20). Lymphocytes were stimulated for 5 hr at 37°C with 70 ng/mL Phorbol 12-Myristate 13-Acetate (PMA; Fisher Scientific) and 700 ng/mL ionomycin (MP-Biomedicals) and treated with 5 μg/mL brefeldin A (MP-Biomedicals) (21). Lymphocytes were then surface stained with anti-CD4 (RM4-5), anti-CD8α (53-6.7), and for exclusion of non-relevant cells, anti-CD45R (B220; RA3-6B2), anti-CD 11b (MI/70), anti-CD11c (N418), and anti-Ly-6G (Gr-1; RB6-8C5), then washed. Cells were then fixed in BD Cytofix/Cytoperm (BD Biosciences) according to manufacturer’s instructions. Next cells were incubated for 45 min at 4°C in 0.5% saponin/PBS with anti-IL-2 (JES6-5H4) and anti-IFNγ (XMG1.2).

### T cell stimulations and Phospho-flow cytometry

For all signaling assays, lymphocytes were serum-starved for 1 hr at 37°C before stimulation in complete media without serum. For evaluation of TCR signaling, cells were stimulated with 20 μg/mL anti-CD3ε (145-2C11; Biolegend) and 50 μg/mL anti-IgG (polyclonal, α-Armenian hamster, Jackson ImmunoResearch) for the indicated times (22, 23). For evaluation of IL-2-mediated pSTAT5 signaling, lymphocytes were stimulated for 15 min with recombinant human IL-2 (rhIL-2; NIH AIDS Reagent Program) at the indicated concentrations. For evaluation of IL-15-mediated pSTAT5 signaling, lymphocytes were stimulated for 15 min with recombinant mouse IL-15 (rmIL-15; Peprotech) at the indicated concentrations. Immediately following stimulation cells were fixed with a final concentration of 1.5% methanol-free formaldehyde (Fisher Scientific; Cat. PI28906) for 15 min, and permeabilized with 500 μL of ice-cold methanol per 1×10^6^ cells for 30 min (22, 24). Cells were washed twice with PBS/2%FBS (Omega Scientific) then stained for 60 min in anti-CD4 (RM4-5), anti-CD8α (53-6.7), anti-FoxP3 (FJK-16s), anti-pSTAT5 (Y694; C71E5; Cell Signaling Technologies) or anti-pS6 (S235/236; D57.2.2E; Cell Signaling Technologies), and, for exclusion of non-relevant cells, anti-CD45R (B220; RA3-6B2), anti-CD11b (MI/70), anti-CD11c (N418), and anti-Ly-6G (Gr-1; RB6-8C5).

### Flow Cytometry Gating Strategies

For analysis of all panels, leukocytes were initially gated on forward and side scatter, then for viability by exclusion of eFluor506 viability dye. For T cell analysis, non-relevant cells were then excluded as described and CD4 or CD8 T cells were gated for feature characterization. Hematopoietic cell populations were analyzed by the indicated surface expression following exclusion of lineage positive cells - LSK (CD127^−^ Sca-1^+^ c-Kit^+^), CLP (CD127^+^ Sca-1^int^ c-Kit^int^), MEP (CD 127’ Sca-1^−^ c-Kit^+^ CD34^−^ CD16/32^−^), CMP (CD127^−^ Sca-1^−^ c-Kit^+^ CD34^+^ CD16/32^−^), and GMP (CD127^−^ Sca-1^−^ c-Kit^+^ CD34^+^ CD16/32^+^). RBC precursors in the BM were analyzed by expression of CD71 and Ter119 following exclusion of CD71^−^Ter119^−^ cells - RBC I (CD71^+^ Ter119^−^), RBC II (CD71^+^ Ter119^+^), RBC III (Ter119^+^ CD71^int^), RBC IV (Ter119^+^ CD71^−^).

### Principal Component Analysis (PCA)

PCA was performed using singular value decomposition in RStudio using R version 3.4.0 through the Bioconductor “pcaMethods” package (Supplemental Methods). PCA was initially performed on 14 parameters acquired from CBC and RBC Ab detection. Regression tree analysis via Chi-square automatic interaction detection, performed using XLSTAT-Base Version 19.4.46344, identified four parameters that contributed the most to the variation, and these four parameters - WBC, RBC, hematocrit, and mean platelet volume - were used for subsequent principal component analyses.

### Statistics

All statistics were performed in GraphPad Prism Version 7 or RStudio with R version 3.4.0. For evaluations based on *a priori* hypotheses, unpaired, two-tailed student’s t tests were performed with Bejamini-Hochberg alpha correction when doing more than 10 tests for a given hypothesis. Bejamini-Hochberg was chosen for alpha corrections as this has stronger statistical power than Bonferroni correction (25). When standard deviations were significantly different between groups, Welch’s correction was applied to the student’s t test. Grubb’s test for outliers was utilized to determine whether significant outliers were present. If significant outliers were present, those data were excluded from analysis. Log-rank (Mantel-Cox) test was performed to compare Kaplan-Meier survival curves. Pearson’s correlations were used to assess linear regressions for peripheral blood data.

## Results

### Early RBC characteristics predict disease kinetics

To assess general survival and disease progression in BALB/c IL-2Rα-KO mice, survival of IL-2Rα-KO and littermate controls was monitored. IL-2Rα-KO mice died due to anemia-induced hypoxia from 18 to 79 days, with a median survival of 26 days (Figure 1A). Twenty-six percent of IL-2Rα-KO mice died between 19 and 21 days of age and 38% died after 30 days. These data suggested that IL-2Rα-KO mice might fall into distinct early and late disease categories based on the age of death.

**Figure 1.**
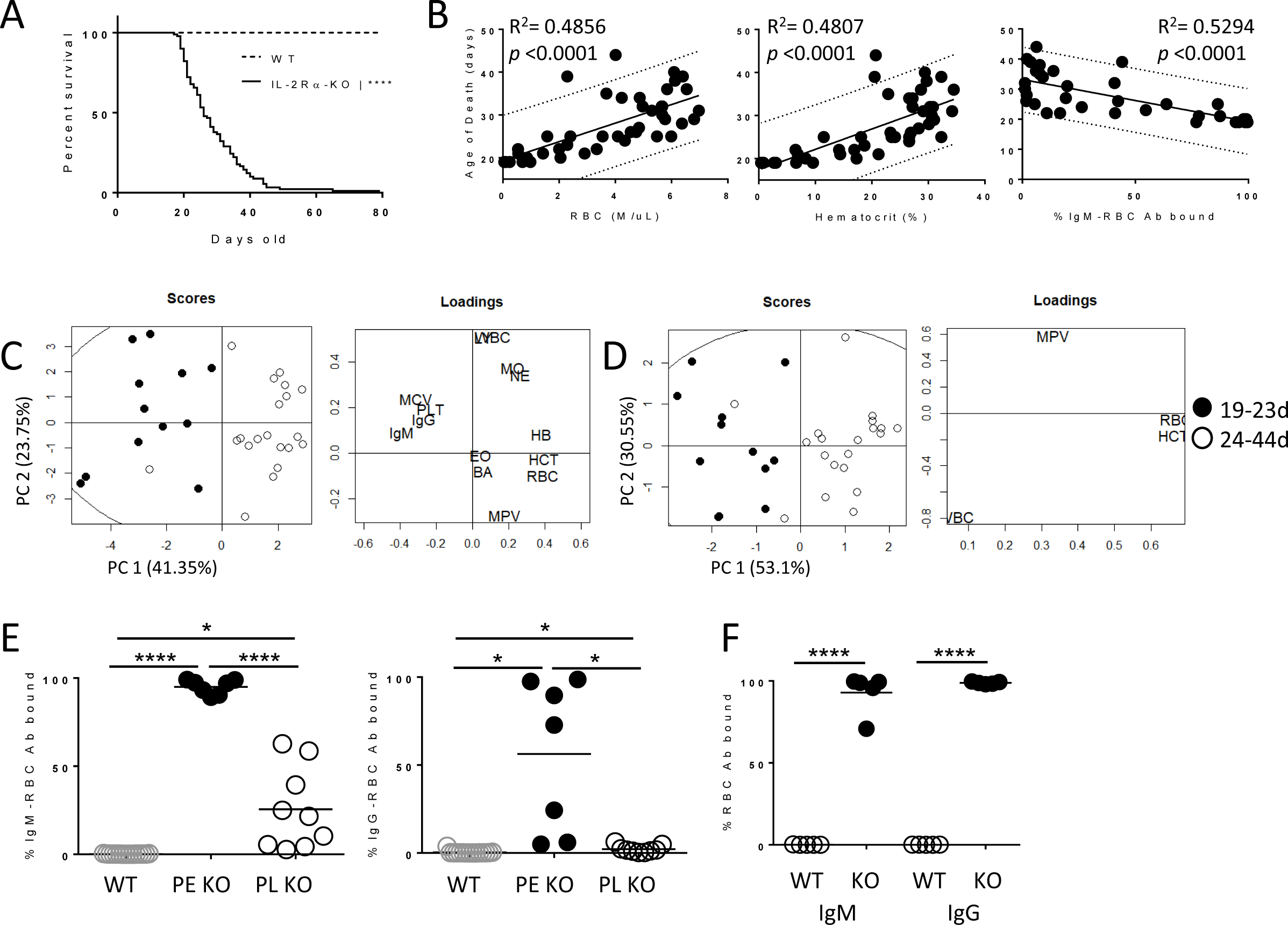
Early RBC characteristics predict IL-2Rα-KO disease kinetics. A) Kaplan-Meier survival curves for WT and IL-2Rα-KO mice monitored for survival from 10 days of age. Statistics: log-rank (Mantel-Cox) test, **** *p* < 0.0001; *n* = 90 IL-2Rα-KO and 47 WT mice. B) Dot plots and linear correlations for data collected from CBC and RBC ab binding were tested against the age of death of IL-2Rα-KO mice. Pearson correlations with the highest R^2^ values are shown. Dotted lines indicate the 95% prediction interval. Statistics: *p* values shown on graphs indicate whether the slope of the linear regression is significantly non-zero; *n* = 44 (RBC and HCT), *n* = 36 (IgM). Principal component analysis by singular value decomposition was performed on peripheral blood collected from 19-day old IL-2Rα-KO using C) 14 variables or D) 4 variables. Percentage on the axes indicate the amount of variation in the data that the given principal component explains. *n* = 32 mice. E) The percentage of RBC bound by IgM or IgG antibodies at day 19. *n* = 7-15 mice per group over >6 independent experiments. F) The percentage of RBC bound by IgM or IgG antibodies of mice age 28 days and over. *n* = 5 mice per group. Statistics: unpaired Student’s t-test with Welch’s correction for variance where appropriate; * p<0.05, **** p<0.0001. Non-significant comparisons are not shown.

To assess disease progression in relation to disease outcome, we sought a biomarker to predict the age of death of IL-2Rα-KO mice. Peripheral blood was collected from IL-2Rα-KO and WT mice at 19 days of age and evaluated for CBC and bound RBC antibodies. Mice were monitored for survival, and the age of death was recorded and correlated to the parameters from CBC and RBC antibody detection. The strongest of these Pearson correlations were RBC concentration (R^2^=0.4856), hematocrit (R^2^=0.4807), and percentage of RBC bound by IgM antibodies (R^2^=0.5294) (Figure 1B). While these showed clear correlations, the 95% prediction interval was ±10 days, indicating these variables alone do not allow for accurate predictions.

Since tests of linear correlations were not on their own strong enough to predict the age of death but did suggest that information in the blood was strongly correlated with the disease outcome, we sought to improve the prediction strength by combining the variables together. PCA was performed on the variables from the CBC and RBC antibody detection. Initial PCA was performed on a total of 14 variables collected from the blood, and these variables separated 96.9% of mice into two groups by age of death with 65.1% (PC1 plus PC2) of the variation explained within the first two principal components (Figure 1C). Regression tree analysis on these data revealed four variables - RBC concentration, hematocrit, white blood cell numbers, and mean platelet volume - as being most important in separating mice by age of death (data not shown). PCA performed using only data from these four variables split IL-2Rα-KO mice into two groups by age of death, a group of mice that died early (between 19-24 days) and a group of mice that died late (24 days or older) with 83.65% of variation explained within the first two principal components (Figure 1D). Using the four blood parameters, the range of death could be predicted in 93.8% of mice at 19 days of age. Overall our results demonstrate that the disease outcome of IL-2Rα-KO mice could be determined and predicted based on blood samples collected early in disease progression.

To assess AIHA disease in relation to disease kinetics, IL-2Rα-KO mice at day 19 were placed into predicted early (PE) or predicted late (PL) disease groups based on the four-parameter PCA analysis. Endpoints in the middle time frame (22-27 days) had higher variability based on the PCA analysis, thus a more stringent breakdown of PE (19-21 days) and PL (28 days and over) disease groups was utilized for disease analysis. RBCs collected from these mice were evaluated for binding of IgM and IgG antibodies. Mice in the PE disease group exhibit a high percentage of RBC bound by antibodies, evidence of severe AIHA (Figure 1E). Mice predicted to die after 28 days of age expressed lower levels of RBC-specific IgM and IgG antibodies at day 19 but did develop higher levels of RBC antibodies at their endpoint (Figure 1E and 1F). This suggests that early AIHA development leads to early death in IL-2Rα-KO mice.

### CD4 and CD8 T cells in IL-2Rα-KO mice exhibit differing activation states

Massive lymphoproliferation and T cell activation is known to occur in BALB/c and C57BL/6 IL-2-KO and C57BL/6 IL-2Rα-KO mice (2, 3), however the preceding events and the immune parameters that correlate with early disease onset in these models are not known. To identify such immune phenotypes, mice were sacrificed on day 19 early in disease onset and separated into their predicted disease PE or PL groups based on the four-parameter PCA and then assessed for T cell numbers and activation by flow cytometry. The ratio of CD4:CD8 in the LN, SPL, and BM of both disease groups showed a shift from the normal ratio of ~2:1 to a 1:1 (Figure 2A). The shift in CD4:CD8 T cell ratio is not a consequence of thymic development, as T cell ratios in the thymus were normal (data not shown). CD8 T cell frequency and total number were elevated in IL-2Rα-KO mice relative to wildtype littermates (Figure 2A and Supplemental Figure 1A). The percentage of CD4 T cells was reduced in the LN and BM but was normal to increased in the SPL (Figure 2A). These data contrast what is known in IL-2-KO mice, as IL-2-KO mice have elevated CD4 and CD8 T cell numbers but maintain normal frequency (1). Comparing PE and PL mice, splenic CD4 and CD8 T cell frequency was higher in PE mice, but these differences were not observed in other organs (LN and BM).

**Figure 2.**
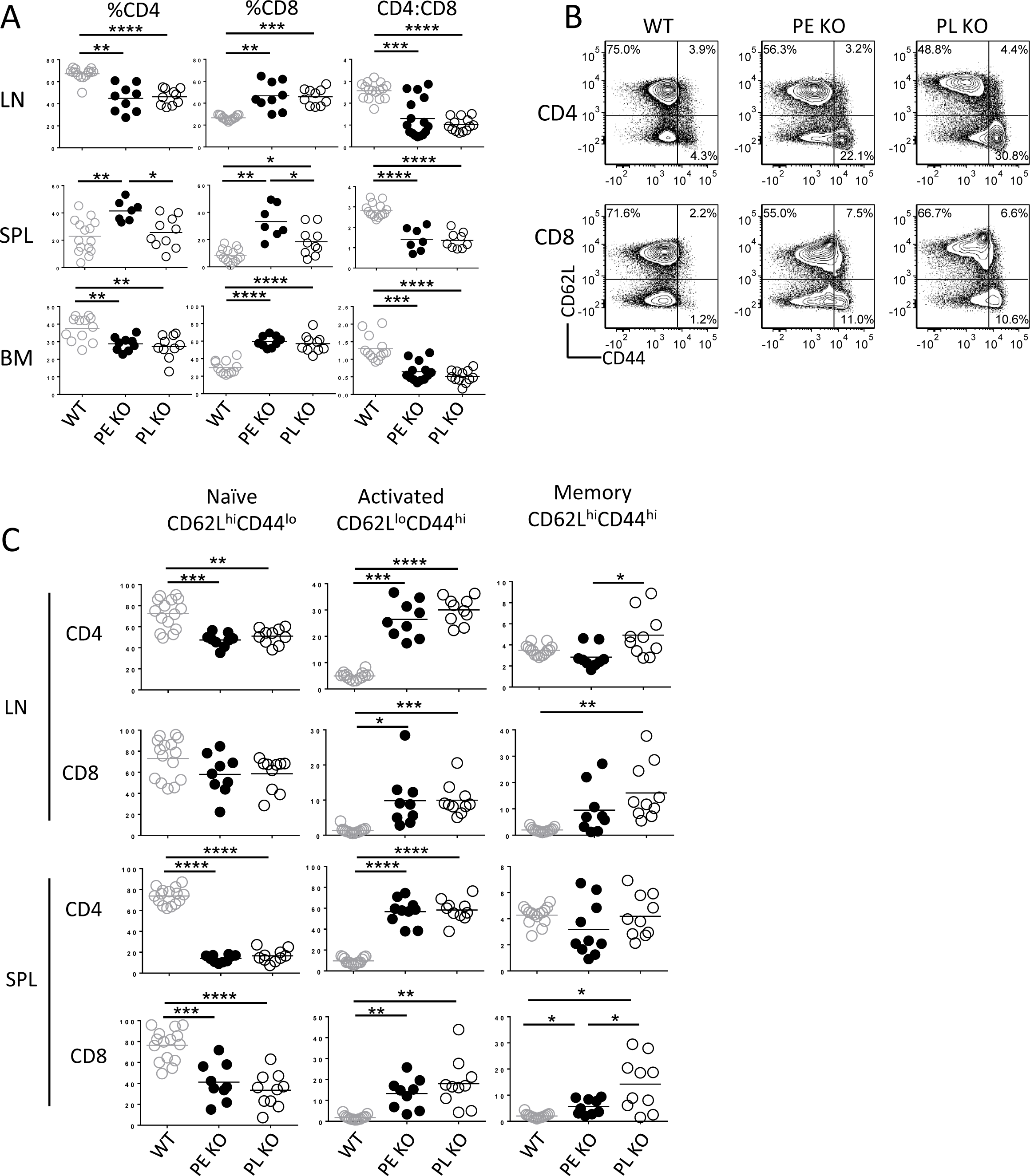
IL-2Rα-KO T cells express markers indicative of developing memory. A) CD4 and CD8 T cell frequency and the ratio of CD4:CD8 T cells in LN, SPL, and BM are shown for WT and IL-2Rα-KO mice. B) Representative flow plots of CD44 and CD62L expression on LN CD4 and CD8 T cells in WT and IL-2Rα-KO mice. C) CD4 and CD8 T cell activation state in WT and IL-2Rα-KO mice. Statistics: unpaired Student’s t-test with Benjamini-Hochberg alpha correction and Welch’s correction for variance where appropriate; * p<0.05, ** p<0.01, *** p <0.001, **** p <0.0001; *n* = 10-16 mice per experimental group in >6 independent experiments. Non-significant comparisons are not shown.

Both PE and PL IL-2Rα-KO disease groups demonstrated increased T cell activation based on reduced CD62L and elevated CD44 expression, as has been observed in IL-2-KO mice (Figure 2B, 2C and Supplemental Figure 1B). In contrast, a higher frequency of IL-2Rα-KO LN CD4 T cells from PL mice exhibited an activation state (CD62L^hi^CD44^hi^) indicative of a developing memory population (Figure 2B and Supplemental Figure 1B). CD8 T cells also exhibited markers indicative of a developing memory population with an increased CD62L^hi^ CD44^hi^ population, and expansion of this CD8 T cell memory population in the SPL associated with late disease kinetics (Figure 2B, 2C and Supplemental Figure 1B). These results indicate an expansion of LN CD4 and splenic CD8 memory cells may correlate with late disease onset.

### IL-2Rα-KO mice exhibit kinetic-dependent differences in CLP and RBC precursor frequency

IL-2-KO mice develop severe BMF contributing to anemia development (5, 26), so we next evaluated whether there were differences in BM populations between PE and PL IL-2Rα-KO mice. Since IL-2Rα-KO mice die from hypoxia, loss of RBCs is critical to disease endpoint. To evaluate the production of RBCs in the BM, unlysed BM from WT and IL-2Rα-KO mice were evaluated for RBC precursor populations using Ter119 and CD71. At day 19, neither predicted disease group showed significant defects in the development and maturation of RBCs (Figure 3A, and Supplemental Figure 3A), despite a decrease in the megakaryocyte-erythrocyte progenitors (Figure 3E). In contrast to disease in IL-2-KO mice, we observe no alteration in erythrocyte precursors in IL-2Rα-KO BM during early disease, although these changes are found in late stage endpoints (Figure 3B and Supplemental Figure 2B).

**Figure 3.**
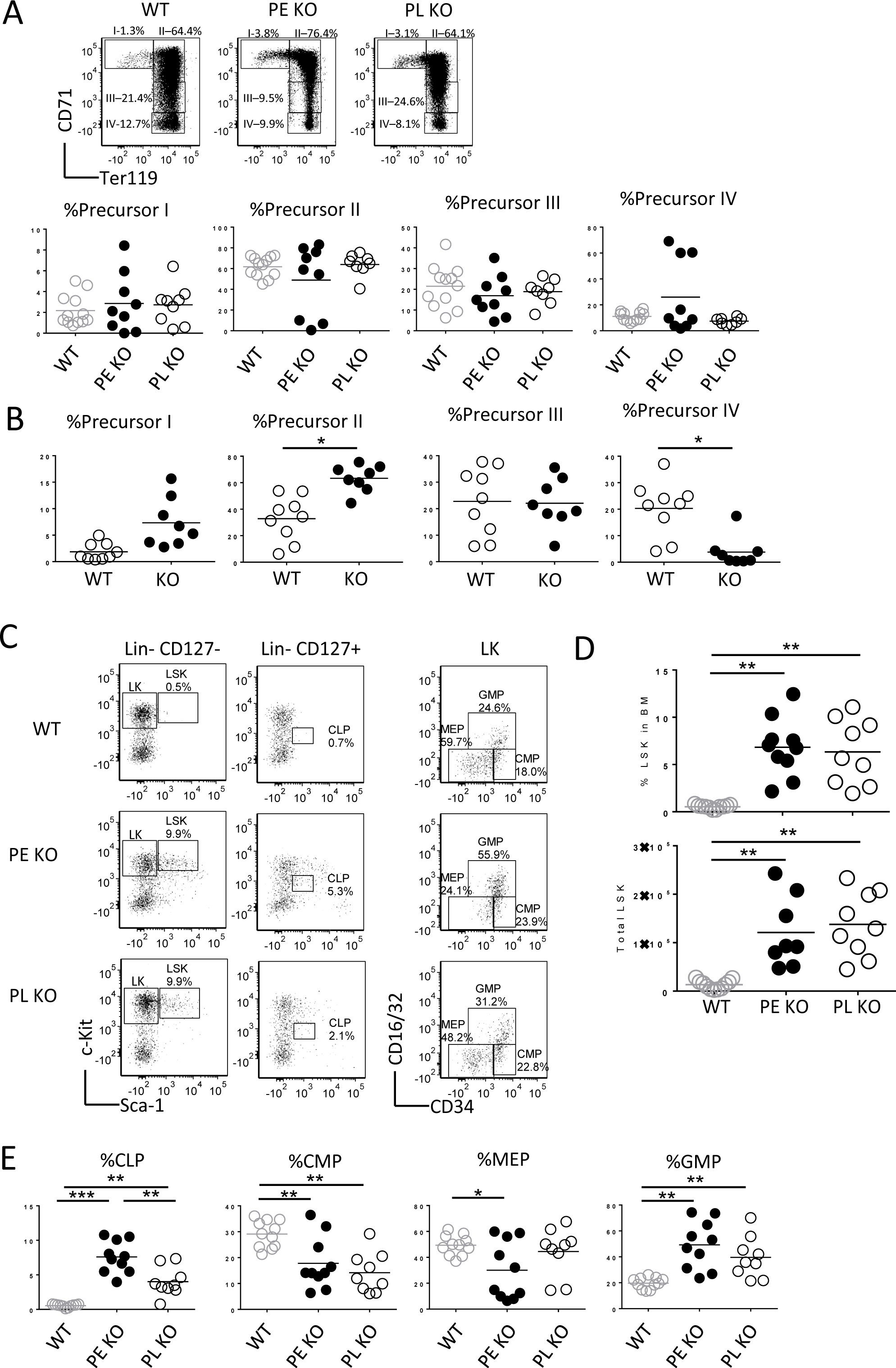
IL-2Rα-KO mice exhibit kinetic-dependent differences in CLP and RBC precursor frequency. A) Representative flow plots and frequency of RBC precursor analysis by expression of CD71 and Ter119 in WT and IL-2Rα-KO mice at day 19. B) Frequency of RBC precursor populations in WT and IL-2Rα-KO mice at day 28. C) Representative flow plots for hematopoietic stem cells and progenitor populations in the BM of WT and IL-2Rα-KO mice. D) Frequency and total number of hematopoietic stem cells in WT and IL-2Rα-KO mice. E) Frequency of hematopoietic progenitor populations in WT and IL-2Rα-KO mice are shown. Statistics: unpaired Student’s t-test with Benjamini-Hochberg alpha correction and Welch’s correction for variance where appropriate; * p<0.05, ** p<0.01, *** p<0.001, **** p<0.0001; *n* = 9-12 mice per experimental group with >6 independent experiments. Non-significant comparisons are not shown.

Next, BM from PE and PL IL-2Rα-KO and WT mice were evaluated for hematopoietic progenitors (Figure 3C, D). IL-2Rα-KO mice in both groups exhibit an increase in the frequency and total number of LSK and CLPs, but a decrease in common myeloid progenitors (CMP) and megakaryocyte-erythrocyte progenitors (MEP) compared to WT (Figure 3D and E and Supplemental Figure 2C). Granulocyte monocyte progenitor (GMP) frequency, but not total number, was increased compared to WT (Figure 3E and Supplemental Figure 2C). Similar to IL-2-KO mice, BM progenitor frequency is altered in IL-2Rα-KO mice. Interestingly, CLP frequency was higher in PE versus PL mice, perhaps leading to the increase in CD4 and CD8 T cells in the SPL of these mice.

### IL-2Rα-KO CD8 T cells express two-fold higher levels of IL-2Rβ than WT CD8 T cells

It is known that a major defect in IL-2-KO and other IL-2 signaling deficient mice is a reduction in the frequency and function of CD4 Tregs, so we sought to assess whether disease outcome was related to Treg frequency at 19 days (4, 27–29). IL-2Rα-KO mice and littermate controls were evaluated for Tregs presence in the LN and SPL. As expected, LN Treg frequency, but not total number, was about half that of WT, but no difference in Treg frequency or number was observed based on disease kinetics (Figure 4A) (4). Surprisingly, splenic Treg frequency was fairly normal in IL-2Rα-KO mice, and the total number of Tregs in both the LN and SPL was significantly increased from WT irrespective of disease grouping.

**Figure 4.**
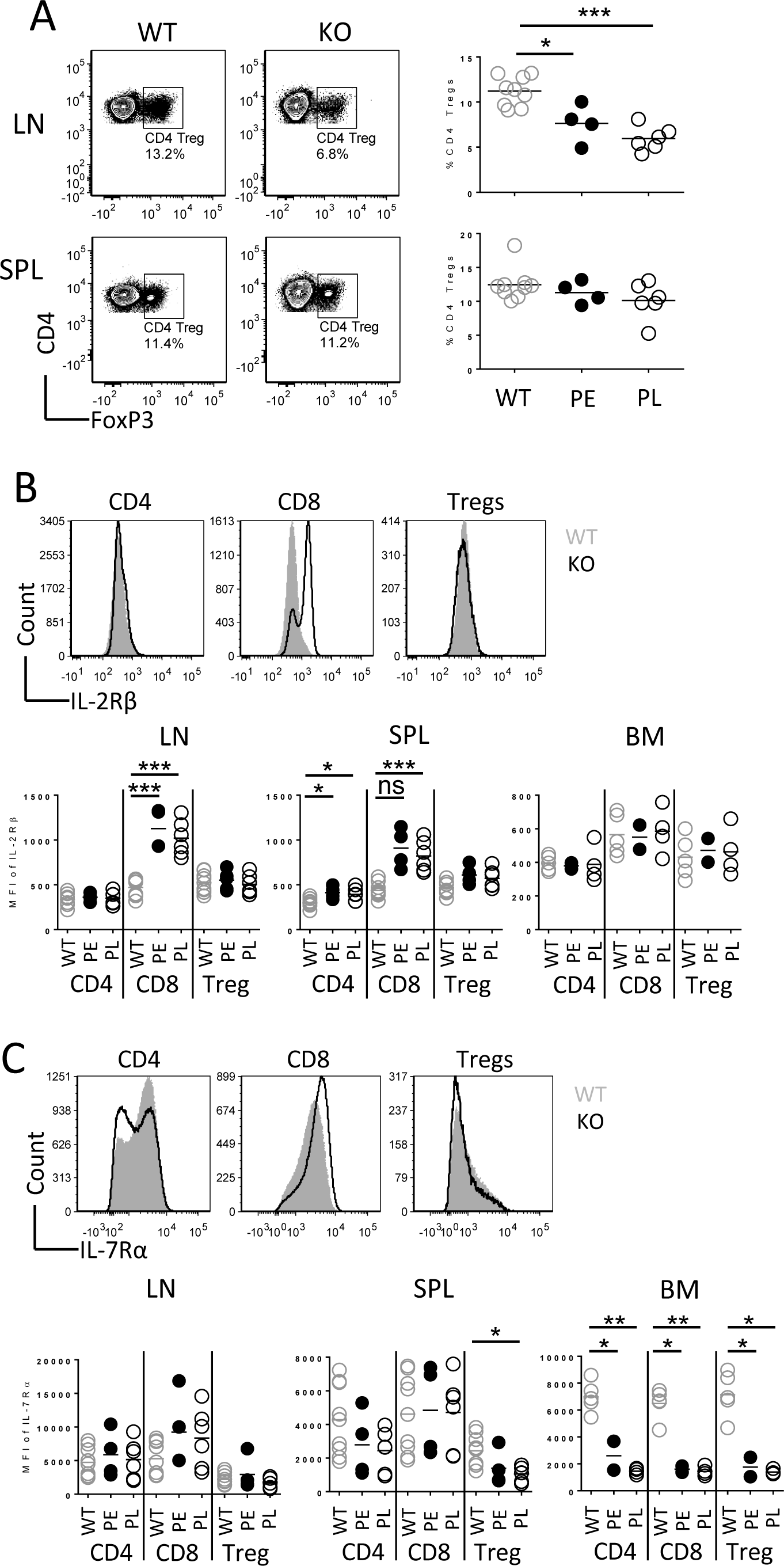
IL-2Rα-KO CD8 T cells express two-fold higher levels of IL-2Rβ than WT CD8 T cells. A) Representative flow plots and frequency of CD4 Tregs in the LN and SPL in WT and IL-2Rα-KO mice. B) Representative histograms showing the expression of IL-2Rβ and IL-7Rα on CD4, CD8, and Treg cells in LN of WT and IL-2Rα-KO mice. C) Summary of mean fluorescent intensity of IL-2Rβ and IL-7Rα on T cells in the LN, SPL, and BM in WT and IL-2Rα-KO mice. Statistics: unpaired Student’s t-test with Benjamini-Hochberg alpha correction and Welch’s correction for variance where appropriate; * p<0.05, ** p<0.01, *** p<0.001, **** p <0.0001; *n* = 5-10 mice per experimental group over 6 independent experiments. Non-significant comparisons are not shown.

The cytokines IL-2, IL-7, and IL-15 provide necessary survival signals to T cells, and elevated IL-7Rα expression on IL-2-KO CD4 T cells promotes autoimmunity (19, 30–33). To assess whether disease outcome was associated with differences in the capacity of T cells to respond to these cytokines, the expression of IL-2Rβ and IL-7Rα on T cells was evaluated. IL-2Rβ expression was elevated on IL-2Rα-KO CD8 T cells but not on CD4 T cells or Tregs in the LN, and was elevated on IL-2Rα-KO CD4 and CD8 T cells in the SPL compared to WT (Figure 4B). IL-7Rα expression was mildly decreased on Tregs in the SPL (Figure 4C). IL-7Rα expression on T cells was drastically decreased in the BM compared to WT, as has been previously observed (Figure 4C) (34). Differences in IL-2Rβ and IL-7Rα expression were observed between CD4 and CD8 T cells in IL-2Rα-KO mice that did not explain differences in disease kinetics. Yet, the cytokine receptor expression levels indicate a differing capacity for CD8 and CD4 T cells to respond to IL-2, IL-7, and IL-15 in the absence of IL-2Rα signaling.

### IL-2Rα-KO T cells are differentially capable of responding to IL-2

We next sought to evaluate whether disease kinetics was determined by production of certain cytokines or differential signaling in response to cytokines. Elevated serum IL-2 levels have been observed in C57BL/6 IL-2Rα-KO mice (35), and IFN_γ_ is known to be critical for early autoimmune disease in IL-2-KO mice (5, 6, 21, 26, 36), so we assessed the capacity of IL-2Rα-KO T cells to produce IL-2 and IFN_γ_. A significant frequency of IL-2Rα-KO CD4 and CD8 T cells produced IL-2, IFN_γ_, and a combination of both cytokines (Figure 5).

**Figure 5.**
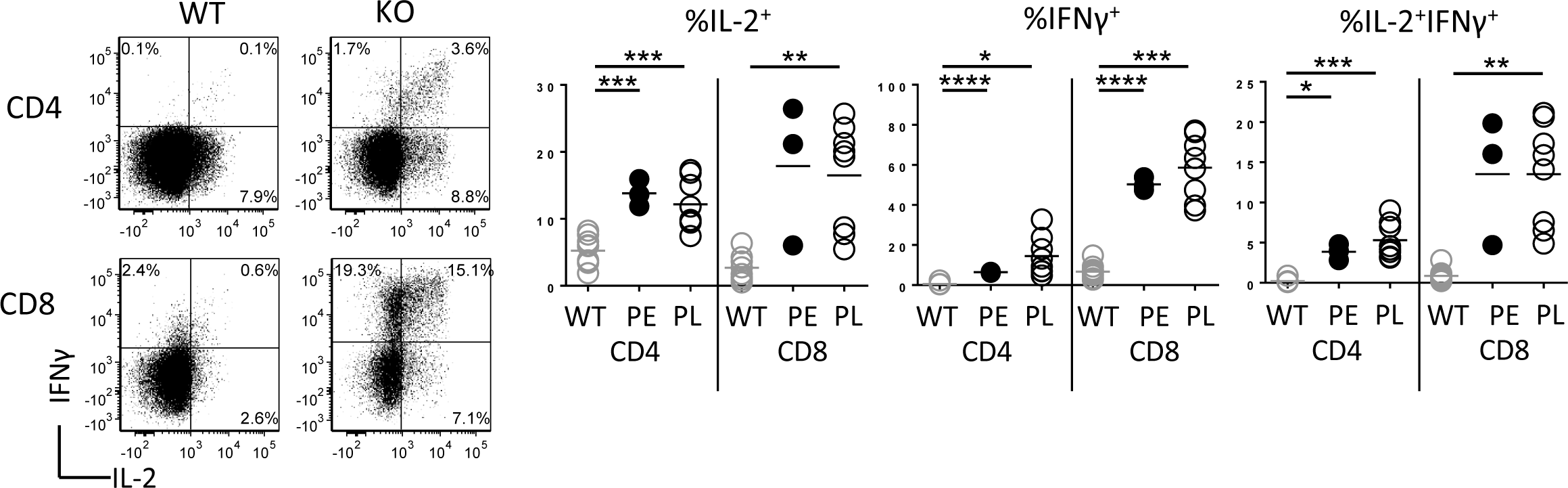
IL-2Rα-KO T cells have increased capacity to produce IL-2 and IFN_γ_. Representative flow plots showing production of IL-2 and IFN_γ_ in WT and IL-2Rα-KO LN T cells post PMA and ionomycin stimulation for 5 hours. Frequency of T cells producing IL-2 and/or IFN_γ_. *n* = 3-9 mice per experimental group over 7 independent experiments. Statistics: unpaired Student’s t-test with Benjamini-Hochberg alpha correction and Welch’s correction for variance where appropriate; * p<0.05, ** p<0.01, *** p<0.001, **** p<0.0001. Non-significant comparisons are not shown.

Recent work evaluating IL-2 signaling in T cells found that IL-2 lowers the threshold of TCR stimulation needed for CD8 T cell proliferation, and that higher basal expression of IL-2Rβ on CD8 T cells allows for sustained IL-2 signaling, leading CD8 T cells to proliferate earlier than CD4 T cells (16, 37). Since IL-2Rα-KO CD8 T cells have two-fold higher IL-2Rβ expression than WT CD8 T cells (Figure 4B), we next evaluated the signaling capacity of IL-2Rα-KO T cells. To assess whether the lack of IL-2Rα would impact TCR signaling, we first assessed phosphorylation of ribosomal protein S6 following TCR stimulation. We found that while S6 phosphorylation kinetics were the same for WT and PE IL-2Rα-KO T cells, PL IL-2Rα-KO T cells had altered kinetics. Additionally, IL-2Rα-KO CD4 and CD8 T cells responded significantly less than WT CD4 and CD8 T cells, with a greater defect present in IL-2Rα-KO CD8 T cells (Figure 6A).

**Figure 6.**
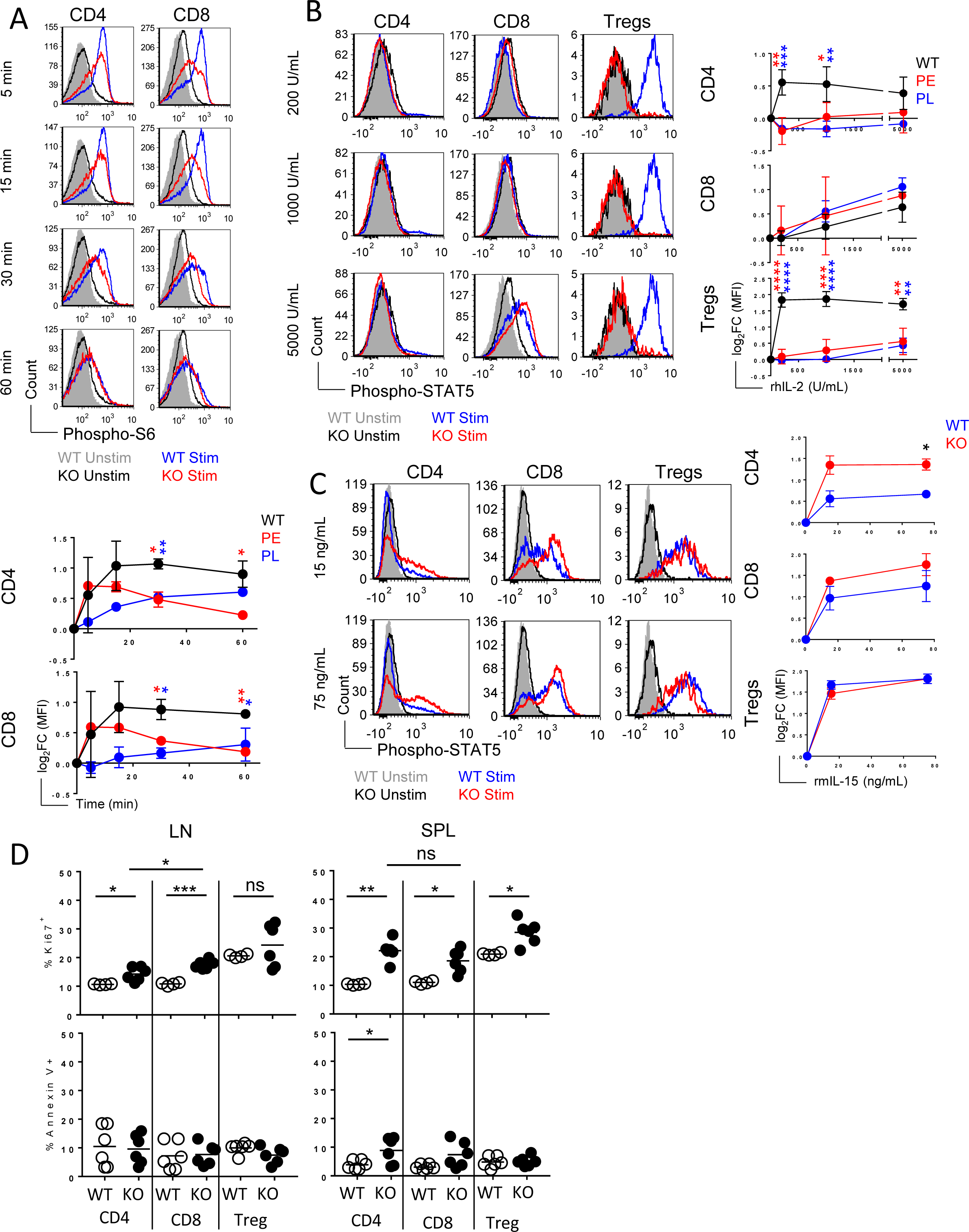
IL-2Rα-KO T cells respond differentially to TCR and IL-2 stimulation. A) Representative histograms of the phosphorylated S6 levels in WT and IL-2Rα-KO LN T cells following TCR stimulation for the indicated time. Log_2_ fold change of S6 phosphorylation over time following TCR stimulation comparing WT versus IL-2Rα-KO mice for CD4 and CD8 T cells. *n* = 2 mice per experimental group over 3 independent experiments. B) Cells were stimulated with rhIL-2 for 15 minutes with the indicated dosage. Representative histograms of the phosphorylated STAT5 levels in WT and IL-2Rα-KO LN T cells following IL-2 stimulation. *n* = 5-7 mice per experimental group over 6 independent experiments. C) Cells were stimulated with rmIL-15 for 15 minutes with the indicated dosage. Representative histograms of the phosphorylated STAT5 levels in WT and IL-2Rα-KO SPL T cells following IL-15 stimulation. *n* = 2 mice per experimental group over 2 independent experiments. D) Frequency of Ki67 positive or annexin V positive WT and IL-2Rα-KO T cells from the LN and SPL are shown. *n* = 6 mice per experimental group over 4 independent experiments. Statistics: unpaired Student’s t-test with Benjamini-Hochberg alpha correction and Welch’s correction for variance where appropriate; * p<0.05, ** p<0.01, *** p<0.001, **** p<0.0001. Non-significant comparisons are not shown.

To assess the ability of IL-2Rα-KO T cells to respond to IL-2, cells were stimulated with recombinant IL-2 at a low (200 U/mL), medium (1000 U/mL), and high dose (5000 U/mL) and STAT5 phosphorylation levels measured. IL-2Rα-KO CD4 T and Treg cells were less capable of signaling in response to IL-2 (Figure 6B). However, STAT5 phosphorylation in response to IL-2 was similar between WT and IL-2Rα-KO CD8 T cells at all doses, likely due to increased expression of IL-2Rβ on IL-2Rα-KO cells (Figure 6B). Together these data suggest that IL-2Rα-KO T cells have differing capacity to respond to TCR and IL-2 stimulation both between CD4 and CD8 T cells, and in comparison to WT T cells, independently of disease kinetics. Further, this differential signaling capacity may contribute to the shift in CD4:CD8 ratio seen in IL-2Rα-KO mice. Since IL-15 shares most of its receptor subunits with IL-2, we sought to assess whether IL-2Rα-KO T cells respond differentially to IL-15 by measuring STAT5 phosphorylation following stimulation with rIL-15. Only IL-2Rα-KO CD4 T cells responded elevated STAT5 signaling relative to WT CD4 T cells (Figure 6C).

To evaluate whether the differential signaling capacity seen for IL-2 and IL-15 has an impact on IL-2Rα-KO T cell proliferation, WT and IL-2Rα-KO T cells were assessed for Ki67 expression. Generally, significantly more IL-2Rα-KO T cells were proliferating than WT T cells, based on Ki67 expression (Figure 6D). Additionally, significantly more LN IL-2Rα-KO CD8 T cells were proliferating than IL-2Rα-KO CD4 T cells (Figure 6D). WT and IL-2Rα-KO T cells had similar levels of apoptosis, except between WT and IL-2Rα-KO CD4 T cells in the SPL (Figure 6D). Together these data suggest that IL-2Rα-KO CD8 T cells receive proliferation signals possibly through combined IL-2 and IL-15 signaling, and that these enhanced signals may allow for the greater expansion and proliferation seen in the IL-2Rα-KO CD8 T cells.

## Discussion

IL-2 and IL-2 signaling deficient mice develop spontaneous, systemic autoimmune disease (1–3). Previous work on C57BL/6 IL-2Rα-KO mice found that approximately 25% of mice died early in disease progression from anemia, with the remaining dying later to a combination of autoimmune phenotypes (3). Similarly, we found that BALB/c IL-2Rα-KO mice died from severe anemia, with approximately 25% succumbing early to advanced AIHA and the remaining dying to a combination of AIHA and BMF later in age.

Prior studies on C57BL/6 mice suggest possible changes in the numbers and ratios of T cells. One study noted increased CD4 and CD8 T cell numbers, but normal frequencies, while another observed an increase only in CD8 T cells (3, 35). We identified a shift in T cell frequencies and total numbers corresponding to an increase in CD4 and CD8 T cells, with a greater increase in CD8 T cells, resulting in an altered CD4:CD8 ratio. Both CD4 and CD8 T cells were activated in IL-2Rα-KO mice, and there was a higher frequency of CD4 and CD8 T cells with a CD62L^hi^CD44^hi^ phenotype in predicted late disease, indicative of a developing memory population (38, 39). However, a corresponding increase in the memory marker, IL-7Rα, on CD4 or CD8 T cells was not consistently seen (38). IL-2Rα-KO CD8 T cells, but not CD4 T cells, expressed higher IL-2Rβ than WT T cells. Although IL-2Rβ expression did not correspond to disease kinetics, this pattern is indicative of memory CD8 T cells, which corresponds with the increased CD62L^hi^CD44^hi^ CD8 T cell frequency found in IL-2Rα-KO mice.

IL-2-KO mice develop severe BMF that contributes to the severe anemia in these mice (5, 26). We observed defects in the BM of IL-2Rα-KO mice indicative of BMF, including increased frequencies of HSC, CLP, and GMP, and reduced CMPs. Surprisingly in light of RBC defects in IL-2-KO mice, these defects did not translate into early defects in RBC precursors in IL-2Rα-KO mice, regardless of disease kinetics. Defects in RBC precursors only became apparent in older IL-2Rα-KO mice, suggesting that BMF does not contribute to early anemia or survival rate.

It has been suggested that increased serum IL-2 levels contribute to the increased memory CD8 T cell population seen in IL-2Rα-KO mice (35). Supporting this, we found that a higher frequency of IL-2Rα-KO T cells produce IL-2 than WT T cells, although IL-2 production did not differ with disease kinetics, nor did the elevated IL-2 result in increased Treg survival, at least in the LN. A prior study suggested that Treg responsiveness to IL-15 may allow Tregs to survive in tissues without IL-2 (40). IL-2Rα-KO CD4 Tregs expressed higher IL-2Rβ levels and were less responsive to IL-2, but were as responsive to IL-15 as WT Tregs, which may explain the increased proliferation and frequency of Tregs in the SPL of IL-2Rα-KO mice. Disease in C57BL/6 IL-2Rα-KO and BALB/c IL-2-KO mice is IFN_γ_ mediated, and T cells from these mice produce significant IFN_γ_ (5, 6, 41). IL-2Rα-KO T cells also produce IFN_γ_, but the frequency of CD4 and CD8 T cells producing this cytokine was not indicative of disease kinetics. However, elevated IL-2 and IFN_γ_ production in IL-2Rα-KO mice may allow for increased or sustained signaling through these receptors.

Recent work found that IL-2 can reduce the threshold of TCR signaling needed to induce proliferation in CD8 T cells, but not CD4 T cells (16). Further, increased expression of IL-2Rβ on CD8 T cells leads to sustained IL-2 signaling and earlier initiation of proliferation (37). We found that IL-2Rα-KO CD8 T cells were significantly less responsive to TCR signaling than WT CD8 T cells, and that this defect was not as obvious in CD4 T cells. Analysis of STAT5 phosphorylation in IL-2Rα-KO T cells following IL-2 stimulation indicated that while CD4 T cells and Tregs were less responsive to lower IL-2 doses than their WT counterparts, IL-2Rα-KO CD8 T cells were not impaired in their ability to respond to IL-2. IL-2Rα-KO CD8 T cells have two-fold higher expression of IL-2Rβ than WT CD8 T cells, which likely contributes to the higher IL-2 sensitivity and responsiveness, although responsiveness to IL-15 was unchanged. This overall increased cytokine signaling may in turn promote the increased CD8 T cell expansion seen in IL-2Rα-KO CD8 T cells, resulting in the elevated CD8 numbers and frequency in IL-2Rα-KO mice.

## Acknowledgements

The authors thank Roy Hoglund and staff of the UC Merced Department of Animal Research Services for animal husbandry care, Dr. David Gravano and the UC Merced Stem Cell Instrumentation Foundry for critical evaluation of the manuscript and their assistance running flow cytometry and advice with gating schemes, and Dr. Jonathan Boyajian and the HSRI Biostats and Data Support Core for statistical consultation and guidance. We also thank our undergraduate researchers, Karelly Barajas, Kendrick Nguyen, and Nicole Quiroz Lumbe for mouse genotyping and assistance with dissections.

